# A 3D-printed cradle for mouse preclinical MRI with an integrated water heating system

**DOI:** 10.1101/2024.12.29.630663

**Authors:** Romain Gaudin, Jeremy Bernard, Melissa Glatigny, Davide Boido

## Abstract

Functional Magnetic Resonance Imaging (fMRI) of small animals is mainly performed under sedation or anesthesia to avoid movement, which is detrimental to image quality. Heating systems to warm the animals usually rely on airflow or heating blankets or pads with circulating water to comply with MR compatibility requirements. However, these solutions are often suboptimal for small animals like mice scanned at ultra-high magnetic fields with long-bore MR scanners. We designed and built an MR cradle with an integrated water chamber, maximizing the contact surface with the mouse’s body.

This large contact surface helps maintain body temperature without overheating the animal, thus reducing the risk of burns and hyperthermia. Our cradle keeps the mouse’s body temperature stable within the physiological range during an MRI session and fits the bore of a Bruker 17.2T scanner. We share the 3D drawings and all the information needed to replicate the cradle. Our design can be adapted to work on preclinical scanners with similar bore sizes and customized to add stimulation devices.

## Introduction

In preclinical research, general anesthesia is still widely used due to the need to avoid movement for *in vivo* data acquisition. It is known that anesthetics induce a reduction in metabolism, which in some instances leads to altered thermoregulation by causing vasodilation, inhibition, or vasoconstriction, resulting in hypothermia^1^. Previous studies have shown that hypothermia induced during general anesthesia can cause bradycardia, disruption of the circadian rhythm, increased risk of infection, as well as difficulties in post-anesthesia recovery^2,3^. Due to their small size, mice have one of the highest surface-to-volume and surface-to-mass ratios in the animal kingdom, making them particularly sensitive to hypothermia^2,4–6^. When no heat source is used during anesthesia, a mouse’s body temperature drops rapidly. To overcome this problem, many heating systems are available on the market. However, most are not MR compatible, and those commonly used inside MR scanners use water circulation pads or blankets placed under or over the animal, respectively. The small contact surface between the animal and the heating surface requires the heating systems to set a high temperature to keep the mouse warm, increasing the risk of burns. In humans, irreversible skin lesions occur after 6 hours of contact at 44°^7^. Consequently, previous studies^8,9^ suggested that the temperature generated by any heating device in the presence of an animal should not exceed 42°C at its surface. Here, we introduce a 3D-printed MRI mouse cradle, compatible with 17.2T, with an integrated heating system. Once placed in the cradle, the animal is almost entirely enveloped by the heating chamber, thus maximizing the contact surface and limiting the difference between the temperature of the heating system and that of the mouse. Here, we determined the effectiveness of this new device in preventing hypothermia during anesthesia and shared the design to replicate our system.

## Materials and Methods

The cradle was designed with SolidWorks (SolidWorks 2023 SP5, Dassault Systèmes). The drawings were transformed from SLDPRT (SolidWorks project files) into STL files (using SolidWorks) suitable for printing with a 3D printer (Form 3+, Formlabs). It is entirely made of resin (Formlabs Grey Pro Resin®). A heated circulator (TC120, GRANT Instruments) immersed in a distilled water bath heated and pushed water to circulate inside the cradle.

Female C57BL/6 mice (n = 3, 6 months old, Janvier Lab, France) were used. This study was carried out in full compliance with French legislation (APAFIS #47074-2024012609362387) and with the approval of the local animal ethics committee. All animals were housed in groups in ventilated cages in a room with controlled temperature and humidity and a 12:12-h dark/light cycle. They were also given ad libitum access to food and water.

Mice were first anesthetized with 3% isoflurane (Isorane 1000 mg/g, Axience) in an induction box before being placed in the cradle, where isoflurane is reduced to a 1.5-2% maintenance concentration. During the tests, two temperature probes are used: a rectal probe to measure the mouse’s body temperature and a probe positioned between the cradle and the mouse to determine the temperature on the surface of the device. The depth of anesthesia was monitored using a pressure balloon system measuring the animal’s respiratory rate, which was maintained between 60 and 90 breaths/minute (Model 1040, SA Instruments, Stony Brook, USA).

The cradle hosting the animal is then positioned inside the MRI to carry out all the measurements over 30 minutes. Any animal temperature below 30°C or above 41°C will immediately terminate the experiment by stopping isoflurane induction. All measurements were taken from when the cradle was set in the center of the MRI. No MRI acquisition was performed in this time interval.

## Results

The design of this cradle (Fig.1) considered several technical constraints: the MR scanner bore diameter (Bruker, BioSpec 17.2T), the ear-bars fixation system, the arrival of gases, the integrated hot water circuit, the fitting of the MRI antenna, and a grove for optical fiber for visual stimulation.

We first measured the maximum temperature at which the water bath temperature should be set to have a temperature not exceeding 40-42°C on the surface of the cradle.

**Fig. 1.**
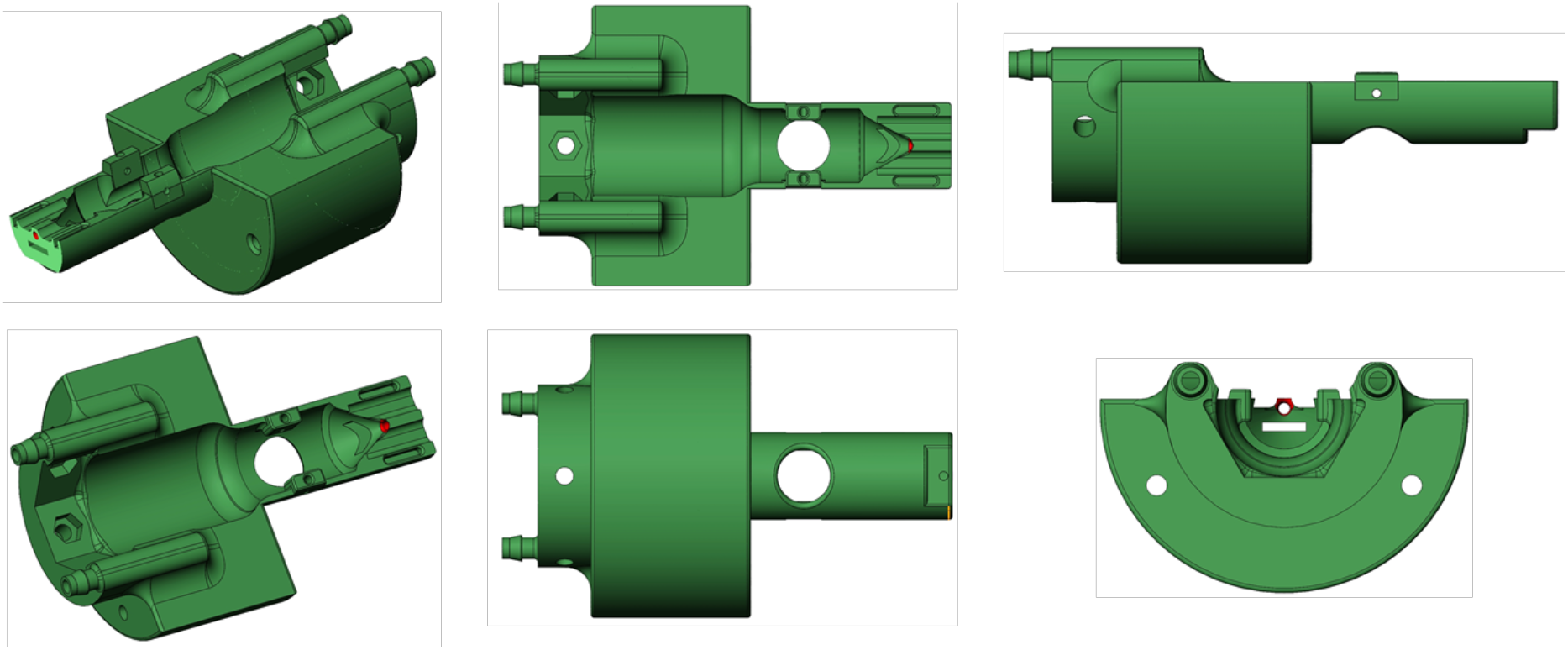
3D extruded design of the warming cradle

The two curves of Figure 2 show that the temperature of the hot water bath and the temperature on the surface of the cradle are highly correlated, with a maximum difference of 10 °C in our experimental conditions. Despite the 8m-long pipe connecting the bath with the cradle (imposed by the ultra-high-field MR scanner), the ∼30 ml water volume inside the cradle has a temperature buffer function. Furthermore, the temperature on the surface of the cradle complies with a safe operational range for animal use.

**Fig. 2.**
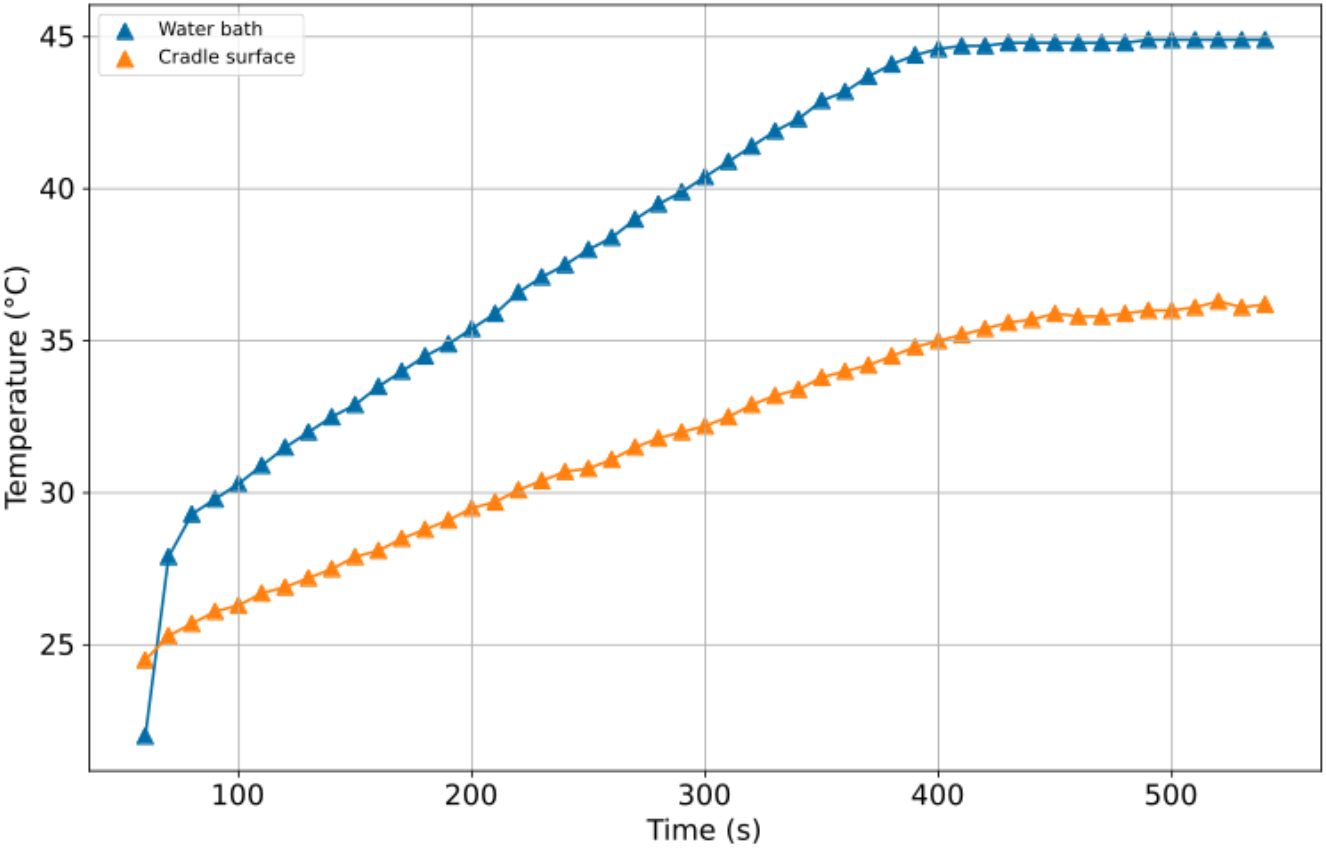
Correlation between water bath temperature and the surface temperature of warming devices.

Then, we tested the new system with a mouse. After falling asleep in the induction box, the animal is placed immediately in the heated cradle, thus avoiding heat loss due to installation time.

The hot water bath was set to 45°C to reach ∼36 °C at the surface of the cradle, equivalent to the mouse’s core temperature (Fig. 3). The bath temperature must be set according to the external environmental temperature, the pipe length, and the mouse’s desired temperature.

**Fig. 3.**
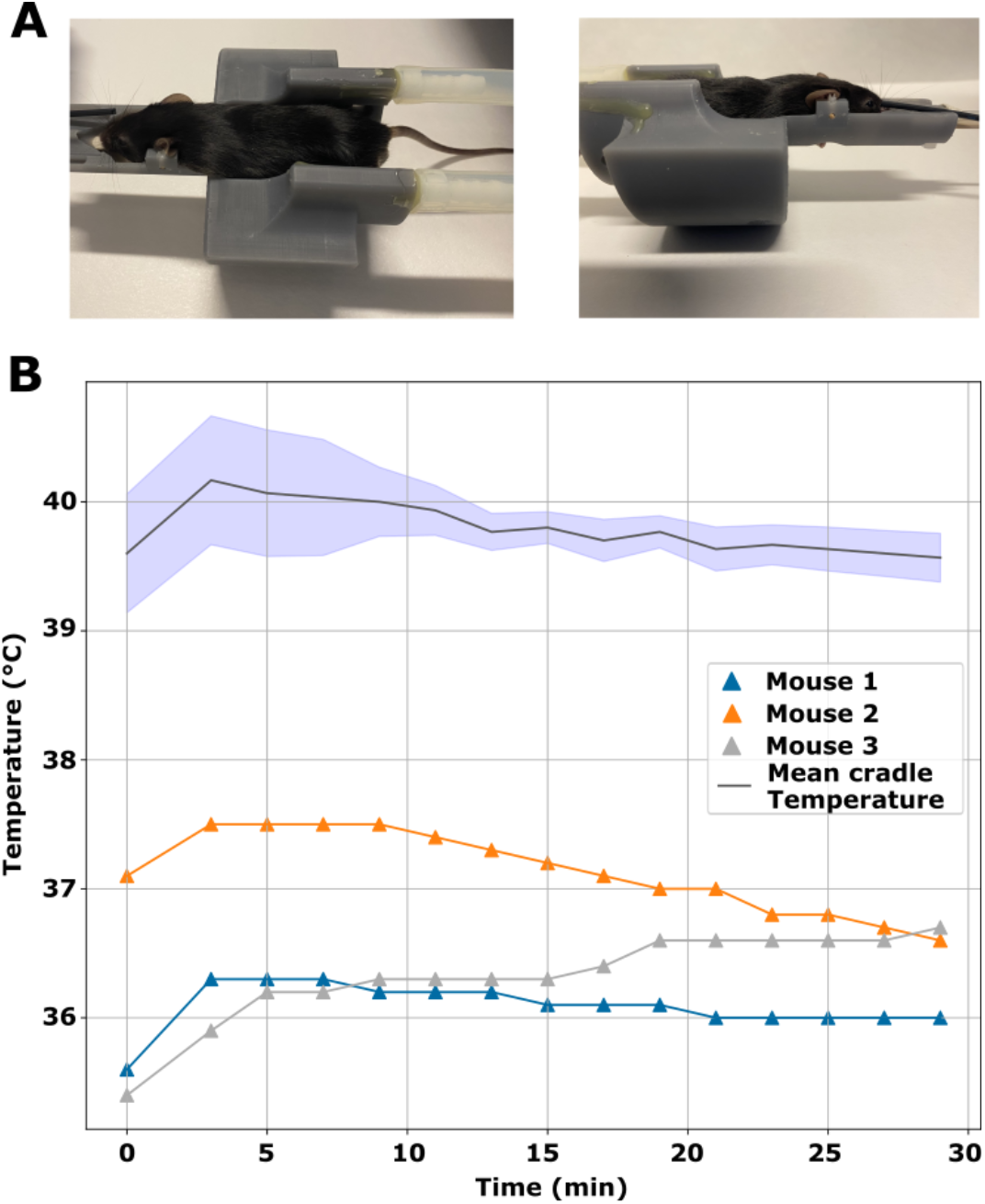
(A) Picture of a mouse in the cradle, with optic fiber for visual stimulation on the right eye (B) Efficacy of temperature maintenance.

An increase of 4 °C in the surface temperature of the cradle was noted with and without the animal (40°C and 36°C respectively) for a hot water bath temperature of 45°C. However, this temperature was still below the recommended threshold of 42°C to avoid any risk of burns. The surface temperature at 40°C set the mouse at a body temperature between 36°C and 36.5°C throughout the 30 minutes of anesthesia by isoflurane inhalation (Fig. 3b).

## Discussion

The state-of-the-art commercially available small-animal cradles for preclinical MR scanners are made in Teflon. However, this material is costly and does not allow much customization of the cradle shape. The new resin 3D printers provide relatively inexpensive cradles (we estimated a cost of €50 per cradle in printable resin), are highly customizable, and have rigidity features comparable to Teflon. However, not all MRI labs have a design and manufacturing facility to conceive and print these cradles. Here, we share detailed information and drawings to produce 3D-printed MR cradles fitting one of our preclinical MR scanners, the Bruker BioSpec 17.2T. This design can accommodate other lower magnetic field scanners with a larger bore. A larger cradle diameter allows for a further increase in the water volume of the inner reservoir, thus increasing the system’s thermal capacity. We showed that, independently of the absolute temperature difference, the cradle surface temperature is highly correlated with the bath temperature over many minutes. This feature will be particularly convenient for researchers working with MRI, relieving the task of temperature checks and tuning usually imposed by less stable warming systems.

Controlling body temperature in mice during general anesthesia is one of the major challenges in preclinical research. Small rodents have a high risk of hypothermia, which causes numerous complications that impact animal physiology, experimental reproducibility, and the quality of the acquired data. This publication is intended to be an ongoing report of the customized MR cradles that we will produce in our laboratory, serving as a unique reference for researchers who will use our systems, eventually with *ad hoc* modifications. Our drawings can be easily modified to accommodate further stimulation devices, such as cannulas and pipes for odor delivery. We hope that any variant of this cradle will be shared with the scientific community.

The drawings and accompanying files of the 3D-printed cradle are available on Zenodo: 10.5281/zenodo.14552887

## Acknowledgment

Financial support was provided by the Agence Nationale de la Recherche (ANR-PRC : CEST-Focus, 2022).

